# Terrestrial Organic Matter Amplifies Methane Emissions Across Sediments of the Mississippi River Headwaters

**DOI:** 10.1101/2024.12.31.630949

**Authors:** Hailey M. Sauer, Leslie A. Day, Trinity L. Hamilton

## Abstract

Terrestrial organic matter (tOM) plays a critical role in aquatic ecosystems, influencing carbon processes and greenhouse gas emissions. Here, we investigate the impact of tOM on methane production in littoral and pelagic sediments from the Mississippi River headwaters using a microcosm approach. Contrary to our expectations, tOM addition universally increased methane production across lentic sediments, with no significant difference between littoral and pelagic zones. Methane production was influenced by select sediment microorganisms, primarily methanogens and lignocellulose degrading bacteria, which responded similarly across different sediment habitats. The study highlights the role of cytochrome-containing methanogens and their syntrophic relationships with fermentative bacteria, emphasizing the significance of microbial community structure in sediment methane dynamics. Our findings suggest that increasing tOM loads to freshwater systems could have broader implications for methane emissions, driven by specific microbial interactions.

**Author Contribution Statement:** HMS and TLH conceived the study and obtained the funds. HMS led fieldwork and microcosm set-up. HMS and LAD analyzed gas samples and HMS performed the data analysis and graphical representation of the results. HMS wrote the first draft of the manuscript, and all authors contributed significantly to the preparation of the final draft.

**Scientific Significance Statement:** As human activities and climate change increase the amount of organic material entering lakes and rivers, understanding the effects this has on greenhouse gas emissions is crucial. Our study reveals that adding terrestrial organic matter to freshwater sediments universally boosts methane production, a potent greenhouse gas. Through the exploration of microbial communities responsible for this process, our research highlights how changes in terrestrial organic matter export to aquatic systems could increase methane emissions from sediments.

**Data Availability Statement:** Additional Supporting Information can be found in the online version of this article, including an extended version of methods and supplementary tables. Sequencing data associated with this paper is available on NCBI, BioProject PRJNA1164797.

## Introduction

Terrestrial organic matter (tOM) plays a significant role in aquatic environments – influencing light, ecosystem metabolism, and food webs (Karlsson et al., 2009, 2012; Pace et al., 2004; Polis et al., 1997). Its role is not only pivotal in shaping the immediate environment but also holds significance in global carbon processes (Guillemette et al., 2017; Heathcote et al., 2015; Lapierre et al., 2013; Tittel et al., 2019; Wilkinson et al., 2013). Current global change scenarios predict an increase in the tOM load to lakes (Intergovernmental Panel On Climate Change (IPCC), 2023). The effects of these changes on lake metabolism and carbon processes are uncertain (Karlsson et al., 2009, 2012; Pace et al., 2004; Polis et al., 1997) but may lead to increases in carbon emissions through terrestrial nutrient subsidies (Lapierre et al., 2013; Tittel et al., 2019). While both carbon dioxide (CO_2_) and methane (CH_4_) can be produced through tOM decomposition in lakes, anoxic CH_4_ production in lake sediments is of particular interest given its higher warming potential (Grasset et al., 2018; IPCC 2023; Lapierre et al., 2013; Tittel et al., 2019).

Methane production by archaeal methanogenesis is strictly anaerobic and occurs predominantly in the sediments where oxygen is limited to the upper millimeters (Buan, 2018; Sobek et al., 2009). In these settings, methanogenic archaea (methanogens) can grow by reducing single carbon compounds (e.g., CO_2_, CO, methanol, methylamines) or acetate through one of several methanogenesis pathways (e.g., hydrogenotrophic, acetoclastic) (Buan, 2018). The use of simplified carbon compounds to fuel this reduction means methanogens are reliant upon other organisms to breakdown more complex organic matter (Liu & Whitman, 2008). As a result, the degradability of the organic matter and the structure of the microbial community equipped to degrade it strongly influences CH_4_ production in sediments (Bastviken, 2009; West et al., 2012).

Several mechanisms influence microbial community assemblage (e.g., environmental filtering and stochastic processes), and within a lake there can be variation in microbial community size and structure – particularly between the littoral and pelagic zones (Cadotte & Tucker, 2017; Nemergut et al., 2011; Niño-García et al., 2016; Vincent et al., 2023). These differences arise in part due to the relative tOM load each of these zones receives (i.e., littoral > pelagic). As a result, CH_4_ efflux from littoral sediments is often higher than that of the pelagic zone (Holgerson & Raymond, 2016). In experimental studies, tOM additions to littoral sediments have led to increases in CH_4_ production contradicting findings that tOM is not a significant contributor to carbon emissions (Grasset et al., 2018; Tittel et al., 2019; Yakimovich et al., 2020). This underscores the significance of OM loads to littoral sediments, while also examining variations in CH_4_ production based on the source, be it terrestrial, phytoplankton, or aquatic macrophytes (Grasset et al., 2018). Considering the varied responses of sediments from a single lake basin to diverse OM sources, sediment regions might react differently due to their unique microbial community composition adapted to most efficiently degrade locally abundant OM compounds. Thus, a focus on sediment from one location leaves unexplored the potential differences in CH_4_ responses to tOM across a lake basin with diverse microbial assemblages.

Here we aimed to assess the spatial heterogeneity of CH_4_ production by pelagic, littoral, and riverine sediments as a response to tOM addition in a hydrologically connected system using a microcosm approach to better understand the biologic mechanisms underpinning observed differences in CH_4_ production in aquatic systems. We expected that (a) CH_4_ production rates would be higher in littoral than pelagic sediments due to a community structure more accustomed to degrading complex tOM, and that (b) variation in CH_4_ production across the system would be related to the abundance and distribution of methanogenic communities.

## Materials and Methods

### Study Site and Microcosm Design

In June of 2020, we collected duplicate gravity cores from five sampling locations within the Mississippi River headwaters system (Fig 1A; Fig S1). Of these locations, two were pelagic, two were littoral, and one was riverine. Itasca State Park (Minnesota, USA) surrounds the system, and the watershed is predominately mature mixed deciduous and coniferous forest. We preformed loss on ignition (LOI) on a homogenized sample from 0-15cm for one core from every sampling location to determine the sediment organic fraction of each site (Table S1) (Dean, Jr., 1974). The remaining core for each site was used to set up microcosms with high, low, and no tOM (i.e., leaf litter) treatment groups. Detailed information regarding sampling locations (Table S1) and the set-up of the microcosms can be found in the supplemental information.

**Figure 1.**
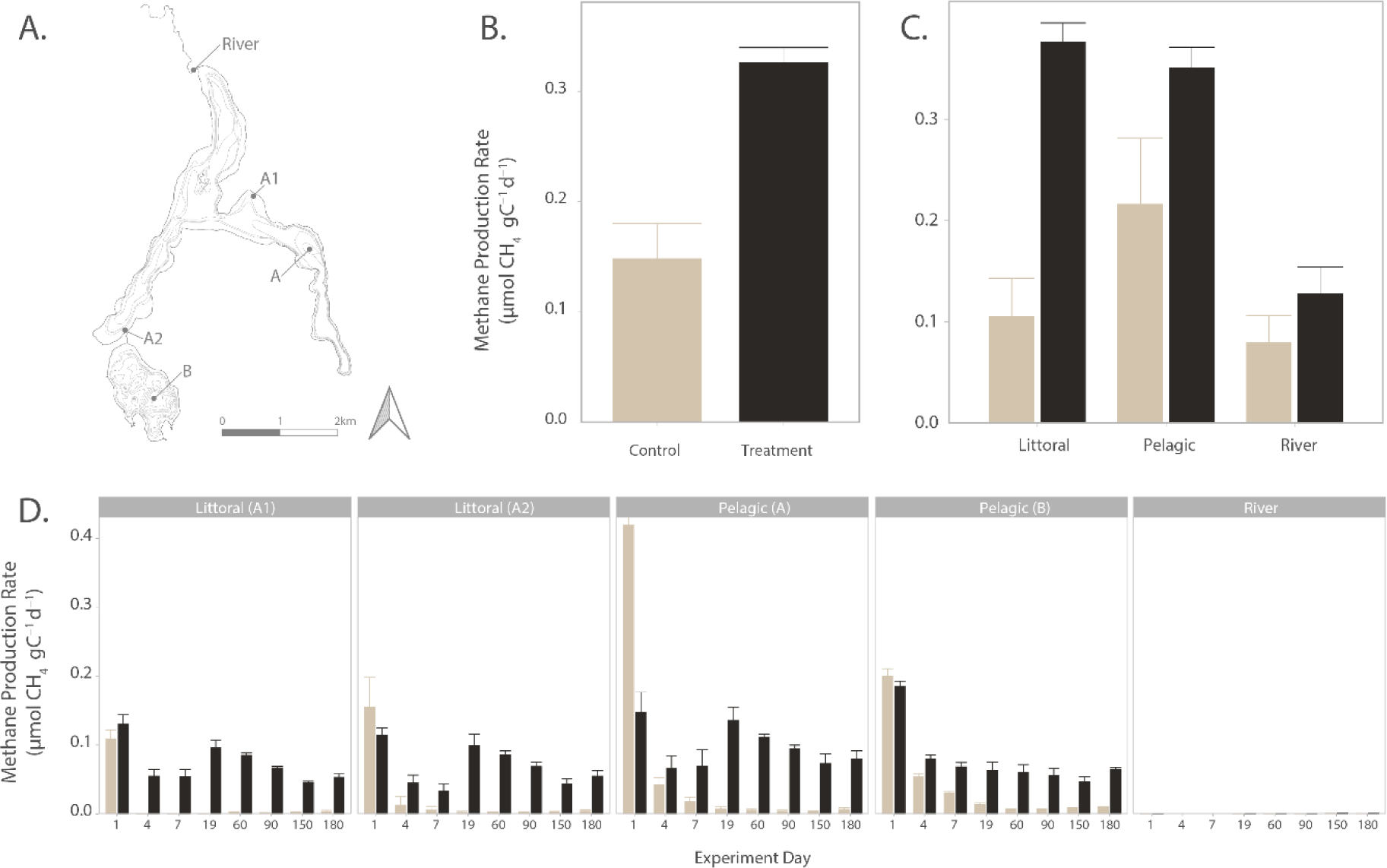
Methane production rates in response to terrestrial organic matter additions across different sediment types in the Mississippi River headwaters. **A)** Bathymetric map (contour 10ft) showing spatial distribution of sampling locations within the Mississippi River headwaters, including littoral (A1, A2), pelagic (A, B), and River sites. **B)** Comparison of CH_4_ production rates (µmol CH₄ gC⁻¹ d⁻¹) between control (no tOM addition) and treatment (with tOM addition) conditions across all sediment types. There was a significant increase in CH_4_ production in the treatment group (Wilcoxon p<0.01). **C)** Observed methane production rates (µmol CH₄ gC⁻¹ d⁻¹) across different sediment types (Littoral, Pelagic, River) under treatment conditions. Both pelagic and littoral sediments exhibited higher methane production rates compared to river sediments, with no significant difference observed between the littoral and pelagic sediments (Kruskal-Wallis & Dunn p<0.01). **D)** Methane production rates (µmol CH₄ gC⁻¹ d⁻¹) for each sediment sampling location over the 180-day incubation period (days 1, 4, 7, 19, 60, 90, 150, 180). The highest production rates were recorded on day 1, particularly in the pelagic A sediment, with rates subsequently decreasing until a secondary peak on day 19 in basin A sediments (Littoral A1, A2, Pelagic A). In all bar plots, the error bars are the standard deviation from the mean for the replicates.

### Carbon to Nitrogen Ratios

We measured total organic carbon (C), total organic nitrogen (N), and isotopic C and N (δ^13^C, δ^15^N) on acidified (1M HCl for 24 hr) sediments pre and post microcosm using a Conflow IV open split interface connected to a Delta V Advantage IRMS (Table S2). We corrected the isotopic values using a set of known external standards including sucrose, GA40, and G41. We exhausted the remaining sample for each site and leaf litter and were unable to determine C and N in 2 samples: Pelagic B high tOM spike and leaf litter.

### Methane Concentrations

During the incubation, we sampled the headspace gas for CH_4_ concentrations at days 1, 4, 7, 19, 60, 90, 150, and 180 using a GC-2014 gas chromatograph. We converted these ppm concentrations to molar concentration using the ideal gas law and Henry’s law, and we normalize production rates per gram C in the microcosm and reported as µmol CH_4_ gC^-1^ d^-1^ (Table S3).

### DNA Isolation, Sequencing, and Post-processing

We isolated DNA from ∼2g of sediment from the initial 0-15cm slurries for each location, ∼6g of leaf litter, and ∼2g of homogenized sediments from the three replicates for each tOM treatment microcosm post experiment. We sent DNA to the University of Minnesota Genomic Center for sequencing and targeted the V3-V4 hypervariable region of the 16S SSU rRNA using primers 341F (5’-CCTAYGGGRBGCASCAG-3’) and 806R (5’-GGACTACNNGGGTATCTAAT-3’) (Yu et al., 2005). We processed the resulting reads using Mothur (v.1.48.0) following the MiSeq SOP (Kozich et al., 2013; Schloss et al., 2009) and aligned our reads using the SILVA database (v.138) (Quast et al., 2013). We performed all subsequent analyses in R, including Latent Dirichlet Allocation (LDA) which correlates microbial communities with relevant environmental factors. A detailed description of LDA is for microbiomes is provided in Sankaran and Holmes (2019) and also outlined in our SI methods (Sankaran & Holmes, 2019).

## Results and Discussion

### Terrestrial OM additions increase methane production in lentic sediments

The addition of tOM to sediments led to a significant increase in the production of CH_4_ across the 180-day experiment in all sampling locations (Fig 1B; Wilcoxon p<0.01). This increase in CH_4_ production came primarily from lentic sediments. There was no difference in CH_4_ production between the pelagic and littoral sediment zones when tOM was added – contradicting our initial hypothesis that littoral sediments would produce more CH_4_. Unamended pelagic sediments produced more CH_4_ than the other unamended sediment types (Fig 1C; Kruskal-Wallis & Dunn p<0.01). Sediments in Pelagic A and Pelagic B both had lower starting C:N ratios (∼8; Table S1) which may explain greater CH_4_ production in the control pelagic sediments. In previous studies, sediment C:N ratios less than 10 lead to greater CH_4_ production regardless of sediment temperature (Duc et al., 2010). Pelagic sediments also receive less oxygen during the open water season and their starting microbial communities may have been more adapted to anoxic microcosm conditions.

Maximum CH_4_ production occurred on day 1 in all lentic sediments (0.85 - 1.85 µmol CH_4_ g^C-1^ d^-1^). While it is possible the peak production on day 1 could be attributed to a rapid, positive priming effect (i.e., mineralization of organic C in the sediments stimulated by the addition of new material), we interpreted the high initial CH_4_ concentrations to be from residual CH_4_ diffused into the headspace during the microcosm setup. Previous research in similar OM amended microcosm studies, using both allochthonous and autochthonous C, support this assumption (Bertolet et al., 2022; Grasset et al., 2018). As a result, the statistics used to compare the overall data, sediment types, and sample sites do not include production from day 1.

After day 1, production decreased until day 19 when there was a secondary peak in basin A sediments (i.e., Littoral A1, Littoral A2, Pelagic A) (Fig 1D). We then saw production decrease until the end of the experiment when there was another, smaller, increase. The non-sinusoidal wave pattern observed in basin A sediments has also been observed in other microcosm studies looking at CH_4_ production response due to tOM additions (Grasset et al., 2018). However, our observed rates are an order of magnitude lower. We attribute this difference to the prior study enriching sediments with nitrogen and phosphorous amended water. When comparing our results with microcosms that solely focus on the sediment microbes reaction to OM (autochthonously produced), we found similar CH_4_ production.

The observed relationship between CH_4_ production in response to tOM with added nutrients, combined with the correlation between trophic status and CH_4_ production, underscores the possible co-limitation in the degradation of tOM and CH_4_ production (Grasset et al., 2018; West et al., 2016). This relationship is further accentuated when we differentiate samples based on their location. Basin A, a meso-eutrophic system, consistently shows higher average methane production across sediment types than basin B. Unfortunately, our study design omitted littoral sediments from basin B. Subsequent experiments could determine whether CH_4_ production patterns (i.e., non-sinusoidal wave) maintain similar characteristics at a basin-level with equal representation of both littoral and pelagic sediments across basins.

Finally, we observed an increase in CH_4_ across all river microcosms, including the control, after 150 days (Fig 1D). While this response may have been a result of the initial river sediments adjusting to the anoxic conditions of the microcosm, we could not confidently determine this. Nevertheless, recent research has highlighted the importance of CH_4_ production in rivers and streams – especially in agriculturally impacted systems (Berberich et al., 2020; Crawford et al., 2016; Stanley et al., 2016). We attributed the low CH_4_ production observed in the riverine treatment condition to a low starting OM percentage (0.3% compared to ∼20% in lentic sediments), and the lack of anthropogenic influence in the protected watershed.

### Sampling location is primary driver of microbial community composition

We recovered 3135 OTUs belonging to 61 phyla across both archaeal and bacterial domains. We classified 67 OTUs as methanogens using taxonomy at the order level (Buan, 2018; Ou et al., 2022; Vanwonterghem et al., 2016). The most abundant OTU in the dataset was a methanogen, and there were 13 others in the top 100 – meaning methanogens are disproportionally abundant relative to their representation (only 2% of the 3135 total OTUs). Four of these OTUs, including the most abundant in the dataset, belong to acetoclastic *Methanosaeta sp.* or *Methanosarcina sp.* in the order Methanosarcinales. Members of Methanosarcinales contain cytochromes and have greater metabolic diversity (i.e., can grow on acetate, methylamines, H_2_) whereas all other known orders cannot use acetate for methanogenesis and are mainly characterized as hydrogenotrophs (using H_2_ and/or formate to reduce CO_2_) (Lyu et al., 2018; Mand & Metcalf, 2019; Thauer et al., 2008).

Most freshwater CH_4_ research focuses on categorizing methanogens based on methanogenesis pathways (i.e., acetoclastic vs. hydrogenotrophic). Here, we considered populations by the presence or absence of cytochromes to overcome limitations caused by multi-phyletic orders for hydrogenotrophic methanogens that differ in energy conservation strategies. By categorizing methanogens this way, we attempted to understand population dynamics and potential nutrient cycling hot spots (Berberich et al., 2020; Biderre-Petit et al., 2019; Tardy et al., 2022). For example, selection for higher affinity to H₂ (non-cytochrome containing) would be advantageous when coexisting with sulfate reducers, which require higher concentrations of electron donors. Alternatively, possessing greater metabolic diversity (cytochrome containing) would facilitate persistence in the environment as available substrates fluctuate. Here, we found methanogens with and without cytochromes across all sediment types and sampling locations and there was no significant interaction between cytochrome status and treatment condition or location. Instead, the methanogen response appears to be a universal increase in abundance regardless of conservation strategy (Fig S2; Table S4). Although our study did not reveal clear niche differentiation based on energy conservation strategy, examining environmental methanogenic communities through the lens of cytochromes is a valuable approach for gaining a deeper understanding of population structure and nutrient dynamics and overcomes limitations and ambiguity inherent to categorizing by methanogenesis pathway.

Finally, and more broadly, we analyzed the structure of the methanogen communities across the sites and treatments using a principal component (PC) analysis (Fig 2A). The first two components explained 75.1% of the variance in the population, which differed based on sediment type (PERMANOVA R^2^ 0.47; p<0.001) and sampling location (PERMANOVA R^2^ 0.64; p<0.001). There is a large range in depth across the four lentic sampling locations (1.10– 27.4m), and this significantly impacted community composition (Pearson R^2^ 0.9868, p<0.001) (Berberich et al., 2020; Ruuskanen et al., 2018; Tardy et al., 2022; J. Zhang et al., 2015). Water column depth correlated with PC2, which explains 23.7% of the variation in community composition. While there were several OTUs that inform this component, there were four OTUs from the genus *Methanospirillum* which were highly abundant in the original sediment community of Basin B and increase in abundance following tOM enrichment – consistent with other experiments in lake and sewage sediments (Chen et al., 2020; Ward & Frea, 1980). In Chen et al. (2020), they found that the abundance of *Methanospirillum* was also significantly correlated with CH_4_ and CO_2_ production; however, the scores from our PC2 did not correlate with methane production rate.

**Figure 2.**
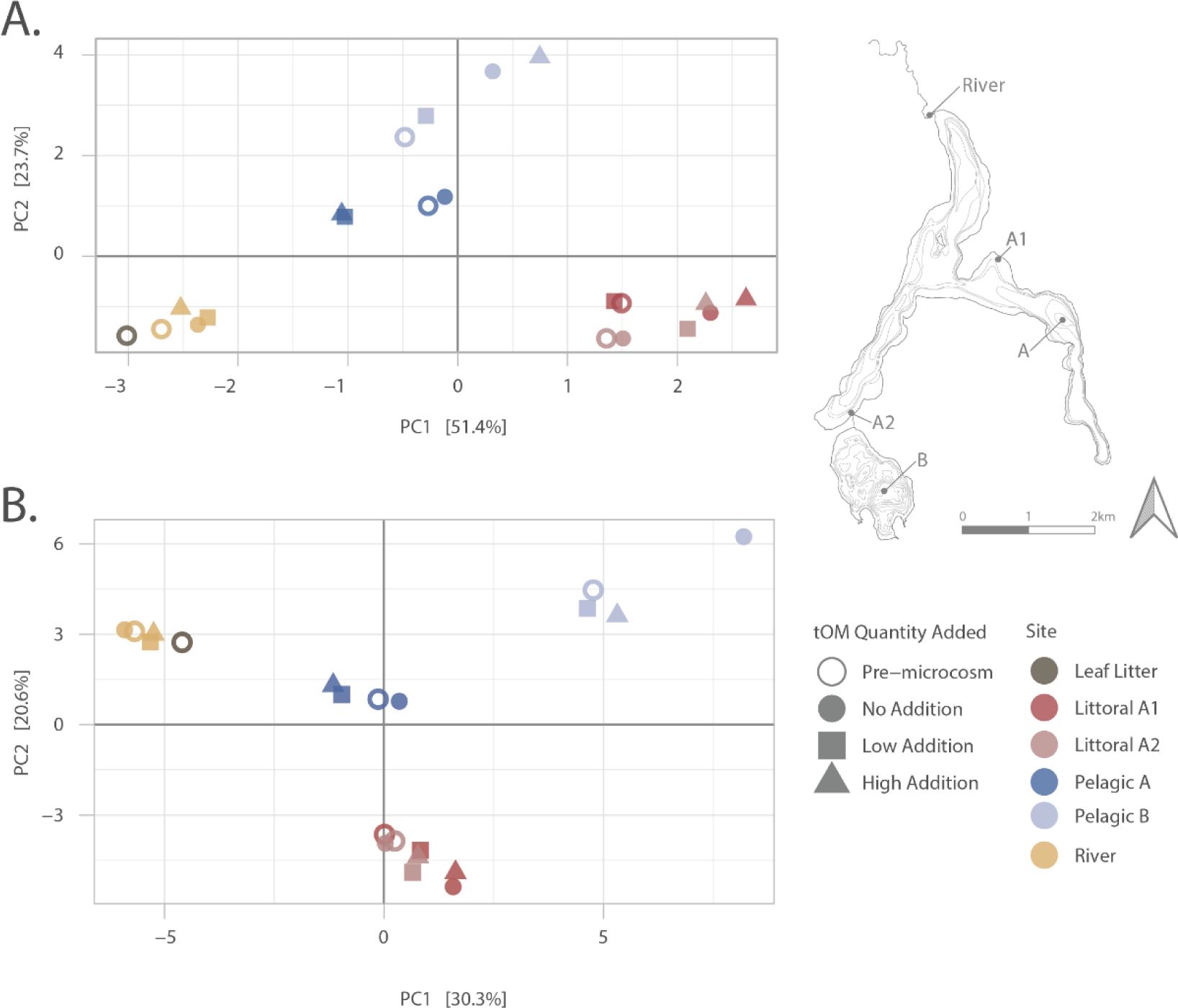
Principal component analysis (PCA) of microbial community composition in response to terrestrial organic matter (tOM) addition across sediment zones. **A)** PCA plot showing the first two principal components (PC1 and PC2) explaining 51.4% and 23.7% of the variance, respectively, in the methanogen community composition of sediments. The points represent different sampling sites (color) and treatment conditions (shape). Methanogen communities are primarily separated by sediment type (PERMANOVA R^2^ 0.47; p<0.001) and sampling location (PERMANOVA R^2^ 0.64; p<0.001). Water column depth strongly correlates with PC2 (Pearson R2 0.9868, p<0.001). **B)** PCA plot showing the first two principal components (PC1 and PC2) explaining 30.3% and 20.6% of the variance, respectively, in the entire microbial community. Sediment type (PERMANOVA R^2^ 0.81, p<0.001) and sampling location (PERMANOVA R^2^ 0.69, p<0.001). The PCA suggest that sediment location is a selecting factor for community assemblage and that water column depth plays an important role in sorting these communities.

Like the methanogen community, the entire bacterial and archaeal community differed based on sediment type (PERMANOVA R^2^ 0.81, p<0.001) and sampling location (PERMANOVA R^2^ 0.69, p<0.001) (Fig 2B). Again, we found water depth to correlate with both PC1 and PC2 (Pearson R^2^ 0.54, 0.83 respectively) and the final C:N ratio from the microcosms was weakly correlated with PC2 (Pearson R^2^ 0.38). However, the mean CH_4_ production for sites and treatments showed no relationship with bacterial composition along either PC axis or final C:N ratio.

### Microbial sub-communities highlight complementary tOM degradation in sediments

Our experimental microcosms showed that while sediment microbial communities differ by sampling location and sediment type, CH_4_ production increased in response to tOM inputs irrespective of community. To further explore the role of unique microbial communities in different sampling locations and sediment types, we used LDA to fit our OTU data into sub-communities and looked for a significant differential abundance in those sub-communities across samples. We found several sub-communities to be differentially abundant when comparing sampling location. On average, sites had 20 significantly different sub-communities. However, low OTU-sub-community probability within made it such that we could not definitively describe patterns, as OTUs often overlap in both significant and insignificant sub-communities. In the pelagic sediments, nearly all sub-community were differentially abundant from river sediments; however, there were no significant sub-communities when comparing the littoral to the pelagic or river sediments.

When we compared these same sub-communities across tOM treatments, we found three (24, 25, & 31) to be differentially abundant given tOM addition (3 levels: Post-No, Post-Yes, Pre; Fig 3A). Given the level order, all results are reported as a change from the Post-No treatment condition. We used a 1% cut off when determining the OTU-probabilities for each sub-community, resulting in a total of 46 OTUs that exhibit significant log2 fold changes across the microcosm treatment. These 46 OTUs are members of eleven different phyla of which the top three are Firmicutes (15 OTUs), Proteobacteria (11 OTUs), and Bacteroidota (9 OTUs). Additionally, we found that between 50-80% of the differentially abundant sub-community OTUs are among the top 100 in terms of percent change (Fig 4). While all three of these sub-communities exhibit a strong and universal response to tOM additions, only sub-community 24 sample probabilities (i.e., gamma) correlated with our observed methane production rates (Pearson R^2^ 0.324) (Fig 3B&3C).

**Figure 3.**
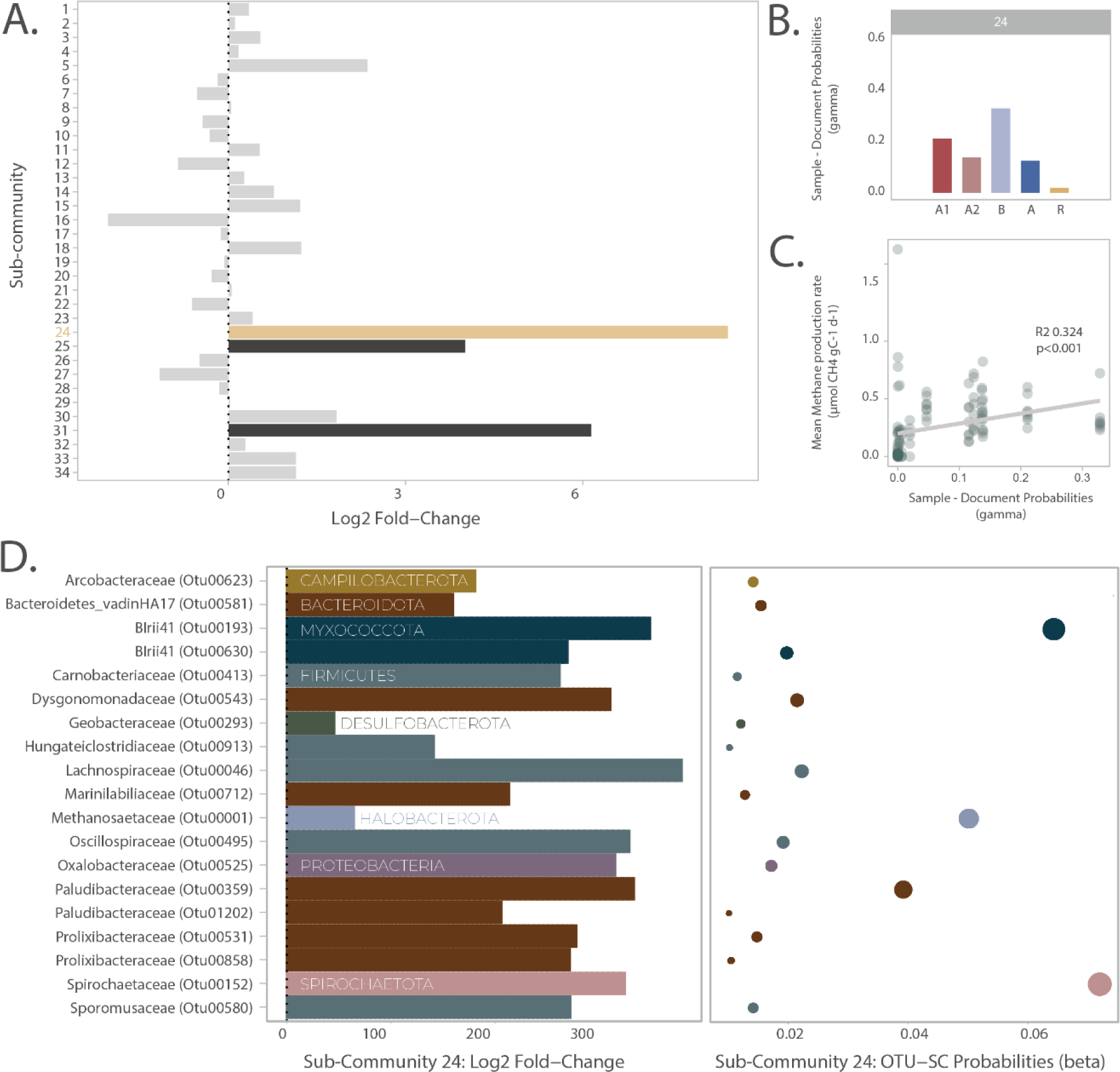
Latent Dirichlet Allocation Model | Microbial Sub-community response to terrestrial organic matter (tOM) addition. **A)** Log2 fold-change in the abundance of 34 identified sub-communities following tOM addition across sediment samples. Sub-communities 24, 25, and 31 showed significant differential abundance in response to tOM addition, with sub-community 24 displaying the most substantial increase (log2 fold-change > 6) and being the only sub-community correlated with increased CH_4_ production. The x-axis represents the magnitude of change, with positive values indicating an increase in abundance post-tOM addition. **B)** Sample probabilities (gamma) for sub-community 24 across the sampling sites. The probabilities reflect the association of sub-community 24 with specific sampling locations, with the highest probabilities observed in Pelagic B and Littoral A1. **C)** Correlation between sample probabilities (gamma) for sub-community 24 and mean CH_4_ production rates (µmol CH₄ gC⁻¹ d⁻¹). There is a significant positive correlation (R² = 0.324, p < 0.001), suggesting that the microorganisms in sub-community 24 may be linked to increased CH_4_ production with respect to tOM addition. **D)** Taxonomic composition and OTU probabilities for sub-community 24. The bar plot on the left shows the log2 fold-change of key operational taxonomic units (OTUs) within sub-community 24, highlighting the predominant phyla, including Bacteroidota, Firmicutes, and Spirochaetota. The bubble plot on the right represents the OTU probabilities (beta) within sub-community 24, with larger bubbles indicating higher probabilities. OTUs such as Treponema sp. (family Spirochaetaceae) and members of the families Prolixibacteraceae and Paludibacteraceae are identified as key contributors to the degradation of recalcitrant organic matter in response to tOM Color in both the left and right graph indicate the phylum of the OTU.

**Figure 4.**
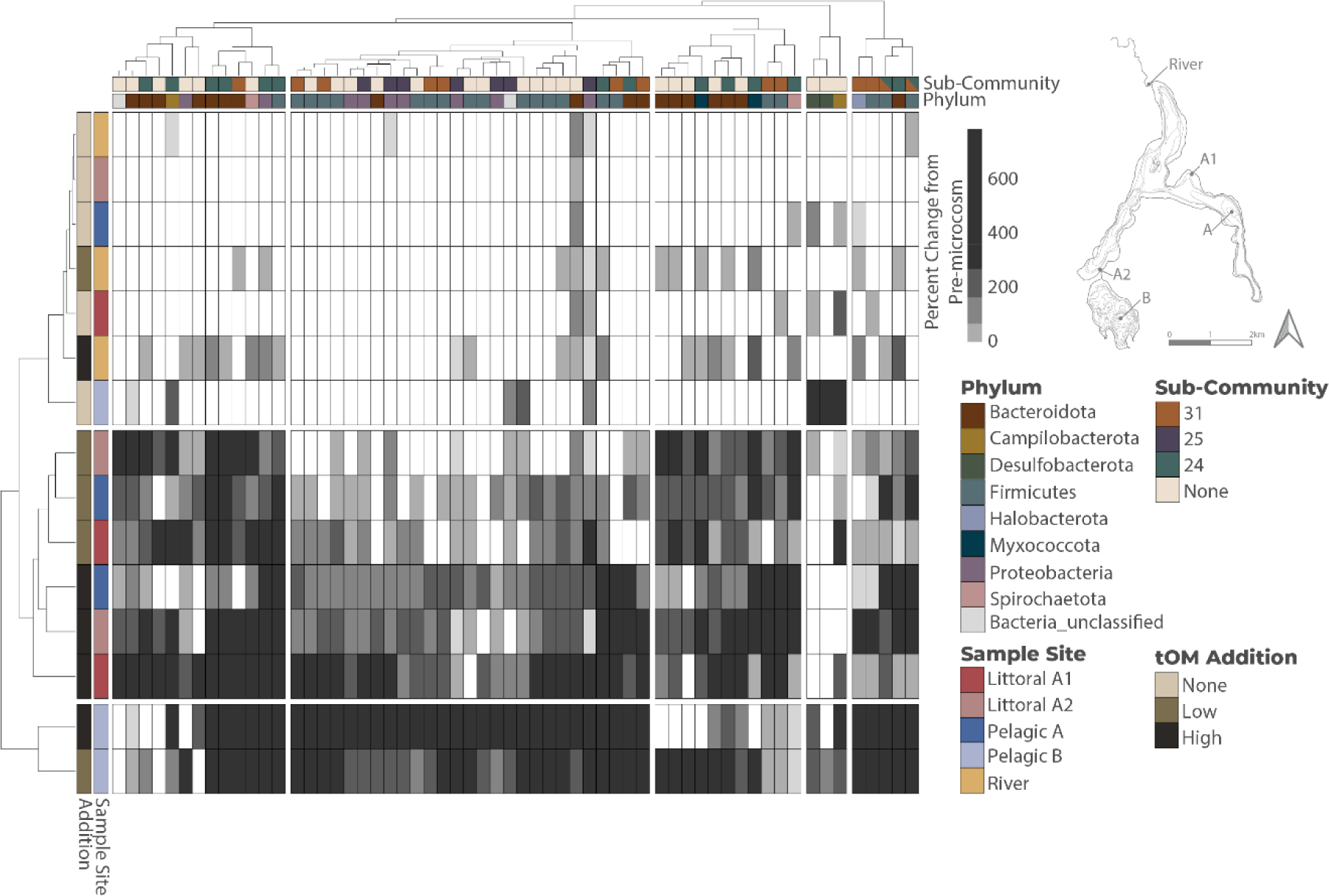
Percent Change of Top 100 OTUs from pre-microcosm to post-microcosm. Heatmap of percent change from pre-microcosm to post-microcosm. Columns represent the top 100 changing OTUs and are Ward D2 clustered. Sub-community status is in the top box wherein the color represents presence within a specific sub-community – split colors indicate multiple sub-community presence. Each OTUs’ phylum is then illustrated, also by color. Rows are Ward D2 clustered samples from the microcosms labeled by their sample location as well as tOM addition. Dendrograms were split in the heatmap at n=3 for the rows and n=5 for the columns based on significant clusters (k-means). Within the top 100 OTUs microcosms cluster primarily by site and tOM status, where similar minimal change occurs in the OTUs with no tOM addition and in the river. Then there is a basin separation between those sites that are in basin A and those in basin B – specifically with respect to increases in OTUs belonging to sub-community 25 (purple) and those clustered with them in basin B.

Sub-community 24 had 19 OTUs and the greatest taxonomic diversity among the 3 differentially abundant sub-communities (8 phyla) (Fig 3D). Many of the OTUs in sub-community 24 were anaerobic microorganisms well adapted to the degradation of recalcitrant organic matter. For example, OTU00152 is a *Treponema* sp. of the family Spirochaetaceae and had the greatest sub-community-probability (∼6%) – meaning it was more frequently associated with the sub-community structure. Members of the *Treponema* genus are noncellulolytic bacteria; however, they aid in the breakdown of cellulosic and lignin materials by interacting with lignocellulolytic bacteria like those in the families Prolixibacteracea and Paludibacteracea, both of which had multiple differentially abundant OTUs in sub-community 24 (Kudo et al., 1987; Leadbeater et al., 2021; Song et al., 2019). Both of these families, like many Bacteroidota, can secrete carbohydrate-active enzymes and have been linked with seasonal patterns of algal detritus degradation in glacial systems, and members of the family Prolixibacteracea specifically have been shown to regulate the decomposition of aquatic plants in salt marshes (Leadbeater et al., 2021; Winkel et al., 2022). The degradation of cellulose in the environment is carried out by a consortia of bacteria that contain complementary enzymes enabling the complete breakdown of bulk tOM (Cragg et al., 2015). The bacterial consortia of sub-community 24 was found at all sampling locations and experienced significant shifts in abundance following the addition of tOM. Within sub-community 24, 15 of the 19 OTUs were among the top 100 shifting OTUs from pre to post microcosm (Fig 4).

In addition to seeing a clear signal response from the fermenting heterotrophic community to tOM treatment, we found two methanogens associated with distinct fermenting community structures – sub-community 24: *Methanosaeta* (OTU 00001) and sub-community 31: *Methanosarcina* (OTU00048). Both methanogens are members of the cytochrome containing order Methanosarciniales and are commonly found in anaerobic digestor sludge where differences in the lipid or polysaccharide content of organic waste can affect the syntrophic partnerships (i.e., two organisms reliant on metabolic cooperation) – leading to different CH_4_ yields (Chang et al., 2018; Kurade et al., 2019; Salama et al., 2019; J. Zhang et al., 2017). In these engineered systems, higher lipid content and syntrophic growth of Firmicutes with *Methanosarcina* results in significantly higher CH_4_ yields (Kurade et al., 2019; Saha et al., 2021). While our sub-community 31 illustrates this syntrophic relationship, the gamma-sample probabilities for sub-community 31 do not correlate with overall CH_4_ production. This was due in part to the unique sample probability for Pelagic B basin with respect to this sub-community, and it potentially highlights that syntroph-methanogen relationships might be lake specific. Despite having low sub-community-OTU probabilities, as result of a relatively low sample size for this analysis (n=21), our LDA was able to uncover commonly occurring community structures like those of anaerobic digestor sludge and weakly correlate a single sub-community’s OTU structure to increased CH_4_ production. To assess the differential abundance of key taxa responsible for CH_4_ production or other environmentally relevant byproducts, future studies could include a larger number of samples.

## Conclusion

The mechanistic relationship between the structure of sediment microbial communities, OM, and CH_4_ production is important to understand contrasting results observed in empirical studies. Here, we found similar patterns of CH_4_ production in response to tOM and found a significant correlation between select hetero-fermentative bacteria in all sediments. We provide an alternative approach to examining methanogen population structures, one in which we consider energy conservation strategies (i.e., cytochrome-containing), and we find that the methanogens of this classification are strongly associated with tOM degrading OTUs in treatment conditions. Ultimately, our findings show that increased terrestrial production and subsequent tOM loads to lake sediments will have implications for lignocellulose degrading bacteria and subsequent CH_4_ production by methanogenic archaea.

## Acknowledgements

HMS and TLH acknowledge funding from NSF grant number 1948058. HMS thanks the Itasca Directors Fellowship for support. LAD thanks the MNDrive (UMII-GA-6798127534) for assistantship We are grateful to K. Fixen, N. Lewis, and J. Eggenberger for instrumentation and lab assistance.

## Supplemental Methods

### Microcosm Set-up

We used the second core from each of the five sites for microcosms – following a 3x3 design with three tOM (dried leaf litter) treatments (0% tOM addition, 5% tOM addition, 15% tOM addition) each with three replicates. We extruded and homogenized the top 15cm of each core under micro-oxic conditions (using an inflatable anaerobic chamber filled with N_2_) and added the homogenized mixture to 125mL serum vials. We filled each vial to a depth of 2cm. For each core we filled the first three vials for the 0% tOM addition treatment and calculated the average sediment weight of the three replicates. We used this value and the starting sediment organic fraction percentage to determine the mass of dried leaf litter needed to increase the organic fraction by 5% and 15%. For tOM additions, we collected and homogenized a dried (105°C for 12h; sieved through No. 5 mesh) mixture of leaves and needles from *Acer*, *Quercus*, and *Pinus* species. We assumed the leaf litter was 100% organic. While there was a statistically significant difference in our tOM addition percentages between treatment (t-test; p<0.001) there was overlap across the percentages and sites, as a result instead of referring to these as firm percentage additions we will refer to them as “low tOM spike” (target 5% tOM) and “high tOM spike” (target 15% tOM). To maintain anoxic conditions, we flushed all the vials with N_2_ before adding sediment and prior to sealing with a gas-tight butyl-rubber stopper and aluminum crimp. Finally, we covered the vials in tinfoil and stored them at 4°C in the dark for the duration of the experiment – 180 days. While we recognize that these conditions do not reflect the *in situ* conditions, our aim was to study methane production dynamics under consistent growth conditions, irrespective of the starting conditions of the initial sediment and abiotic forces.

### Methane Concentrations

We collected 10mL from the headspace of each vial using a gas-tight syringe and injected the sample into Labco Exetainers that were pre-flushed with helium. Exetainers were stored inverted at 4°C until they could be further processed. We replaced the microcosm headspace after every gas pull with 11mL N_2_ gas to maintain headspace pressure and avoid methanogenesis inhibition caused by the accumulation of CH_4_ or CO_2_ (Grasset et al., 2018). We measured CH_4_ concentrations using a GC-2014 gas chromatograph equipped with a flame ionization detector and an 80/100 Porapak N column (6ft x 1/8in x 2.1mm SS). The carrier gas was argon run at a 25mL min^-1^ flow rate, and the calibration standards ranged from 200 to 60,000ppm. We then converted these ppm concentrations to molar concentration using the ideal gas law and Henry’s law. All production rates reported in the text were normalized per gram of C in the microcosm and reported as µmol CH_4_ gC^-1^ d^-1^. We used the values obtained from our total organic carbon analysis to normalize production rates by gram carbon in the supplemental materials. To calculate production rates per gram organic C OUT, we assumed the litter was 80% organic matter with a C:N ratio of 20 (Hornbach et al., 2021). Due to product backorders and shipping delays during the pandemic, we did not have enough Exetainers and were unable to pull gas samples from the Mississippi River site for days 4 and 7.

### DNA Isolation, Sequencing, and Post-processing

During the initial microcosm set up, we subsampled ∼2g of sediment from the 0-15cm slurry and ∼6g of leaf litter and stored these at -20°C until extraction. At the end of the experiment, we combined and homogenized the sediment from the three replicates for each treatment. Again, we collected a ∼2g from the pooled and homogenized sediment and stored it at -20°C until extraction. We extracted triplicate DNA for each sample using ∼0.25g of sediment or leaf litter using a Qiagen Dneasy PowerSoil Pro kit following manufacturer protocols including 4°C incubation steps. We performed negative controls by carrying out extractions on blanks, using only reagents with no sample. We determined the final bulk DNA concentrations using a Qubit dsDNA HS Assay kit and Qubit Fluorometer. We did not detect any DNA in our blanks (detection limit for the assay kit is 10pg/µL). We then pooled 10µL of each DNA yielding replicate and recalculated bulk DNA for the pooled sample. We sent all DNA yielding samples and blanks to the University of Minnesota Genomic Center (UMGC) for sequencing.

The UMGC prepared libraries for the samples for Illumina sequencing using a Nextera XT workflow and 2x300bp chemistry. They targeted the V3-V4 hypervariable region of the bacterial and archaeal 16S SSU rRNA gene using primers 341F (5’-CCTAYGGGRBGCASCAG-3’) and 806R (5’-GGACTACNNGGGTATCTAAT-3’) (Yu et al., 2005). The amplicon preparation performed at the UMGC have been shown to be quantitatively more accurate and qualitatively complete than existing methods (Gohl et al., 2016). We recovered a total of 352,403 raw reads from 24 samples, including blanks and leaf litter. We processed these reads using Mothur (v.1.48.0) following the MiSeq SOP (Kozich et al., 2013; Schloss et al., 2009). We aligned our reads using the SILVA database (v.138) and removed chimeras with vsearch (v2.17.1) (Edgar et al., 2011; Quast et al., 2013). Finally, we classified the sequences as operational taxonomic units (OTUs) using a 97% similarity threshold and assigned taxonomy using the SILVA database (Glassman & Martiny, 2018; Stackebrandt & Goebel, 1994). After processing we had 273,151 reads across the 24 samples.

All further analyses were conducted in R (v4.3.2) using the following packages: tidyverse, phyloseq, vegan, DESeq2, pheatmap, MicrobiomeStat, topicmodels, and ldatuning (Grün & Hornik, 2011; Kolde, 2019; Love et al., 2020; McMurdie & Holmes, 2013; Nikita, 2020; Oksanen et al., 2009; R Core Team, 2018; C. Zhang, 2022). Prior to analyzing the community composition of these sites, we filtered the data by removing any OTU that did not have 2 or more counts in at least 5% of samples. We also removed the nine OTUs which had reads in blank samples, each OTU having a single read. After filtering, the average number of reads per sample was 13,762 and the minimum and maximum read depths were 8,858 and 20,302 respectively. OTU data have a strong positive skew due to many zero counts. To diminish these effects, we used a variance stabilizing transformation (Love et al., 2020). Log-like transformations such as this bring count data to near-normal distributions, produce larger eigengap values, and lead to more consistent correlation estimates all of which influence downstream analysis (Badri et al., n.d.). To compare microbial community structure before and after the microcosm experiment, we conducted a principal component analysis (PCA) of the entire microbial community and the methanogen populations. From both PCAs we pulled the scores of the top two PCs and used those as variables for difference in composition when determining if sediment community composition influences CH_4_ production. We determined methanogens based on taxonomy and compared the methanogen composition based on energy conservation strategies (i.e., those with or without cytochromes) (Buan, 2018; Ou et al., 2022; Thauer et al., 2008). Finally, we took two approaches to compare the change in populations from initial sediments to post-microcosm. First, we calculated the percent change in each OTU (after adding 10% the lowest observed pre- and-post microcosm abundance to any 0 values). We then selected the top 100 OTUs that had the greatest percent change from initial sediment to post-microcosm (eq. 1).

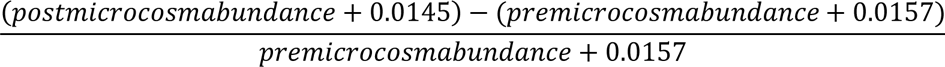

Second, we used Latent Dirichlet Allocation (LDA) or topic modeling to examine the structural differences across microcosm treatments. LDA is a mixture model which correlates microbial communities with relevant environmental factors of interest. The advantages of using LDA over other mixed model approaches is that LDA allows fractional membership allowing samples to be composed of multiple sub-communities. The application of LDA models to microbiome datasets has been described in detail by Sankaran and Holmes 2019 (Sankaran & Holmes, 2019). Briefly, we determined the relevant number of topics or sub-communities (k=34) using the FindTopicsNumber function in ldatuning using a Gibbs sampling method and CaoJuan2009, Arun2010 metrics, and Deveaud2014. Then we conducted the LDA model again using Gibbs sampling and the topicsmodels package. We then converted our LDA model back to a phyloseq object to further assess the differential abundance in the 34 sub-communities across select parameters (e.g., treatment addition, treatment quantity, and sample site). For this, we use the linda function in MicrobiomeStat with an alpha of 0.01, winsorization of outliers at 3%, and zero count data handling set to imputation – where in zero counts are given values with respect to sequencing depth (Zhou et al., 2022). From both approaches we aggregated the list of 100 most changed OTUs and those with >1% OUT-sub-community probabilities in the significantly different sub-communities (n=3; 46 OTUs) and evaluated how the abundances of those OTUs explained methane production rates.

## Supplemental Figures & Tables

**Table S1 | Sampling Site Details**

**Table S2 | Carbon and Nitrogen Data**

**Table S3 | Methane Concentration & Production Rate Calculations**

**Table S4 | Cytochrome Status of Methanogens**

**Figure S1.**
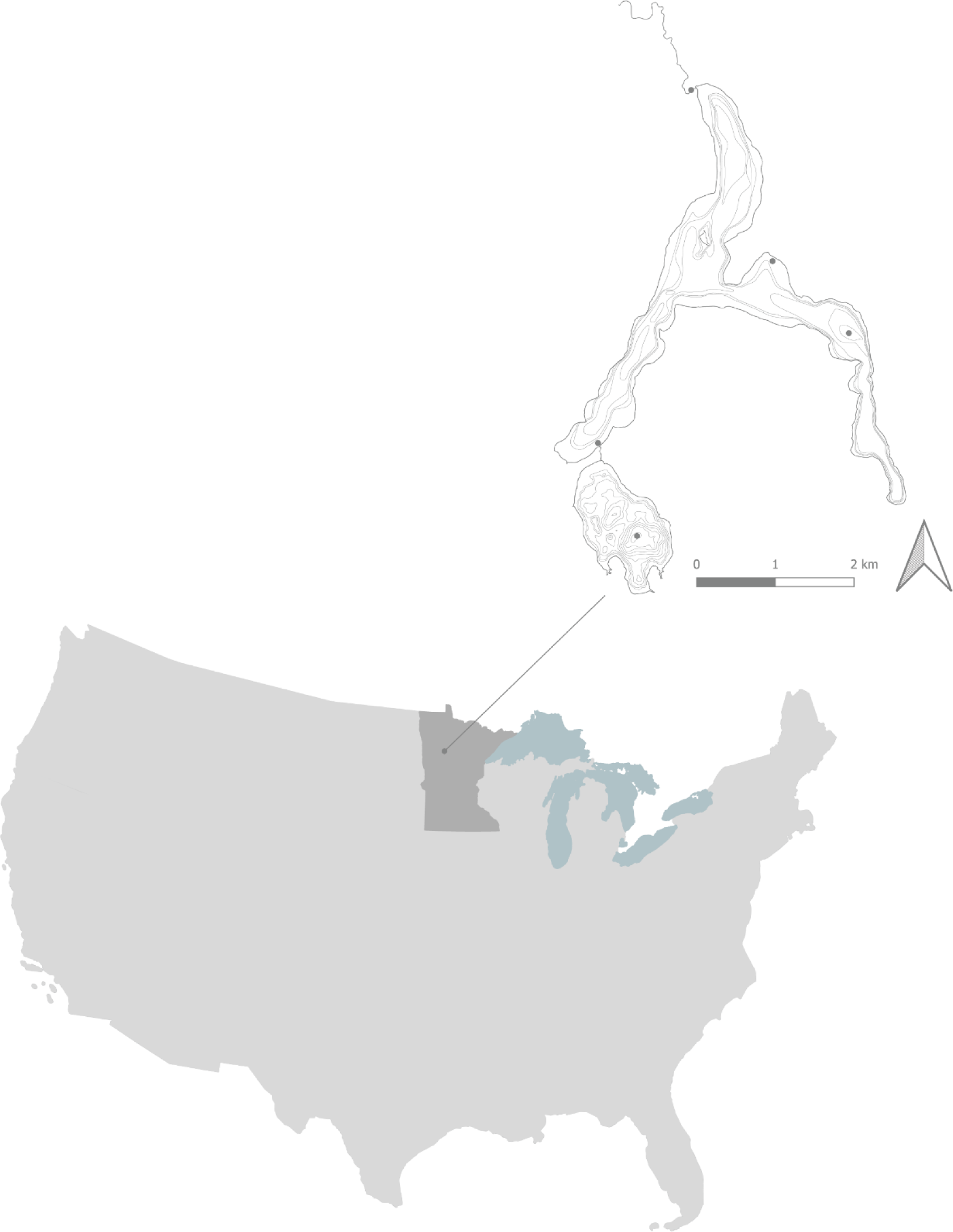
Map. Bathymetric map (contour 10ft) showing spatial distribution of sampling locations within the Mississippi River headwaters, including littoral (A1, A2), pelagic (A, B), and River sites. Scale bar is for the bathymetric map. Elk Lake (basin B) and Lake Itasca (basin A) are the headwaters lakes of the Mississippi River located in Clearwater County, Minnesota USA.

**Figure S2.**
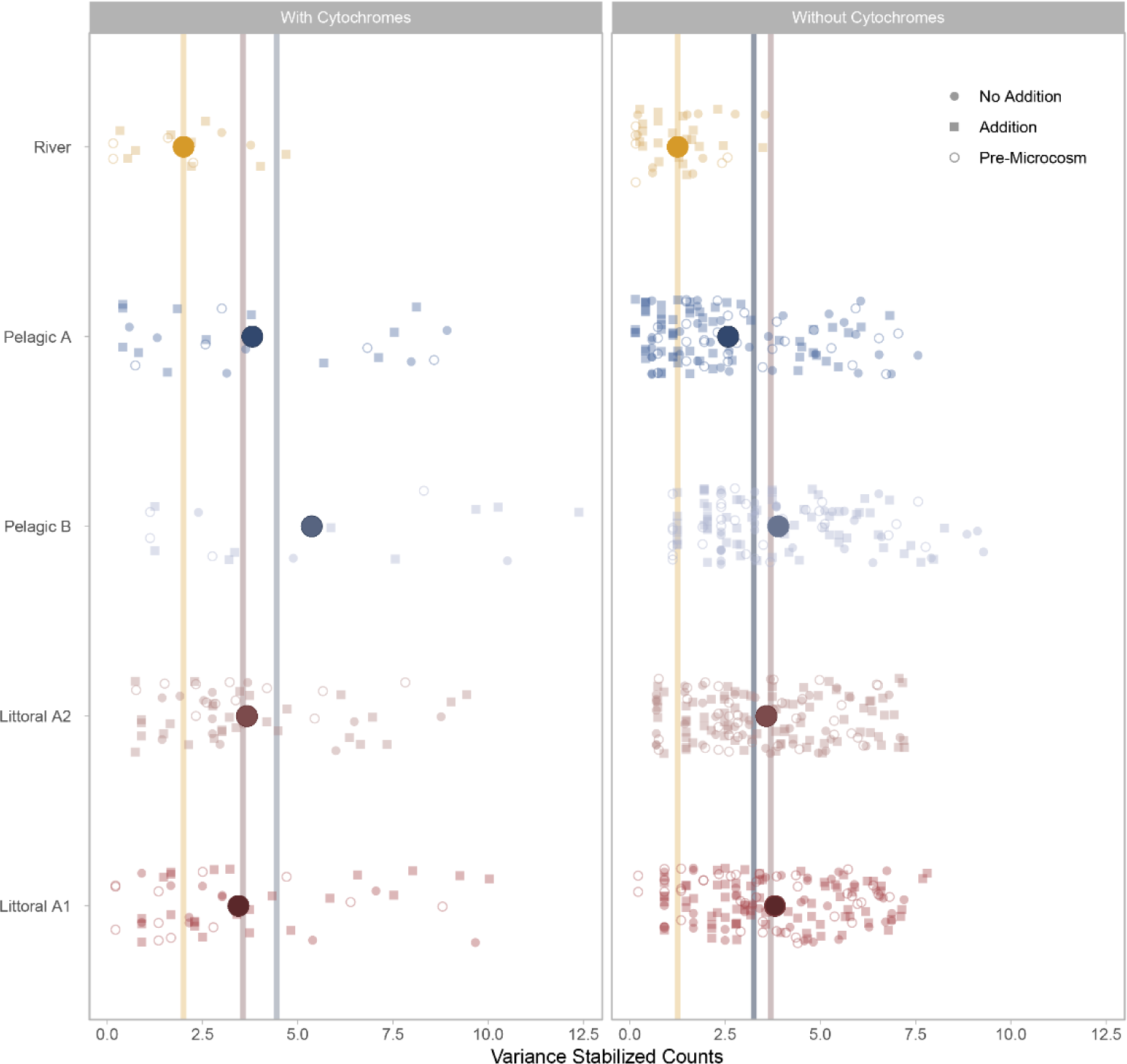
Methanogen Distribution. The distribution of variance stabilized counts of methanogens with cytochromes (left) and without cytochromes (right). Each point is an OTU where color represents location and shape the microcosm status: pre-microcosm (empty circle), no addition (filled circle), and addition (square). Large circles are the average counts per site and lines are the average counts per sediment type: blue (pelagic) and red (littoral).

## References

Badri, M., Kurtz, Z. D., Müller, C. L., & Bonneau, R. (n.d.). Normalization methods for microbial abundance data strongly affect correlation estimates. 21.

Bastviken, D. (2009). Methane. In Encyclopedia of Inland Waters (pp. 783–805). Elsevier. 10.1016/B978-012370626-3.00117-4

Berberich, M. E., Beaulieu, J. J., Hamilton, T. L., Waldo, S., & Buffam, I. (2020). Spatial variability of sediment methane production and methanogen communities within a eutrophic reservoir: Importance of organic matter source and quantity. Limnology and Oceanography, 65(6), 1336– 1358. 10.1002/lno.11392

Bertolet, B. L., Koepfli, C., & Jones, S. E. (2022). Lake Sediment Methane Responses to Organic Matter are Related to Microbial Community Composition in Experimental Microcosms. Frontiers in Environmental Science, 10. https://www.frontiersin.org/article/10.3389/fenvs.2022.834829

Biderre-Petit, C., Taib, N., Gardon, H., Hochart, C., & Debroas, D. (2019). New insights into the pelagic microorganisms involved in the methane cycle in the meromictic Lake Pavin through metagenomics. FEMS Microbiology Ecology, 95(3), fiy183. 10.1093/femsec/fiy183

Buan, N. R. (2018). Methanogens: Pushing the boundaries of biology. Emerging Topics in Life Sciences, 2(4), 629–646. 10.1042/ETLS20180031

Cadotte, M. W., & Tucker, C. M. (2017). Should Environmental Filtering be Abandoned? Trends in Ecology & Evolution, 32(6), 429–437. 10.1016/j.tree.2017.03.004

Chang, S.-E., Saha, S., Kurade, M. B., Salama, E.-S., Chang, S. W., Jang, M., & Jeon, B.-H. (2018). Improvement of acidogenic fermentation using an acclimatized microbiome. International Journal of Hydrogen Energy, 43(49), 22126–22134. 10.1016/j.ijhydene.2018.10.066

Chen, H., Ye, J., Zhou, Y., Wang, Z., Jia, Q., Nie, Y., Li, L., Liu, H., & Benoit, G. (2020). Variations in CH4 and CO2 productions and emissions driven by pollution sources in municipal sewers: An assessment of the role of dissolved organic matter components and microbiota. Environmental Pollution, 263, 114489. 10.1016/j.envpol.2020.114489

Cragg, S. M., Beckham, G. T., Bruce, N. C., Bugg, T. D., Distel, D. L., Dupree, P., Etxabe, A. G., Goodell, B. S., Jellison, J., McGeehan, J. E., McQueen-Mason, S. J., Schnorr, K., Walton, P. H., Watts, J. E., & Zimmer, M. (2015). Lignocellulose degradation mechanisms across the Tree of Life. Current Opinion in Chemical Biology, 29, 108–119. 10.1016/j.cbpa.2015.10.018

Crawford, J. T., Loken, L. C., Stanley, E. H., Stets, E. G., Dornblaser, M. M., & Striegl, R. G. (2016). Basin scale controls on CO2 and CH4 emissions from the Upper Mississippi River. Geophysical Research Letters, 43(5), 1973–1979. 10.1002/2015GL067599

Edgar, R. C., Haas, B. J., Clemente, J. C., Quince, C., & Knight, R. (2011). UCHIME improves sensitivity and speed of chimera detection. Bioinformatics, 27(16), 2194–2200. 10.1093/bioinformatics/btr381

Glassman, S. I., & Martiny, J. B. H. (2018). Broadscale Ecological Patterns Are Robust to Use of Exact Sequence Variants versus Operational Taxonomic Units. mSphere, 3(4). 10.1128/mSphere.00148-18

Gohl, D. M., Vangay, P., Garbe, J., MacLean, A., Hauge, A., Becker, A., Gould, T. J., Clayton, J. B., Johnson, T. J., Hunter, R., Knights, D., & Beckman, K. B. (2016). Systematic improvement of amplicon marker gene methods for increased accuracy in microbiome studies. Nature Biotechnology, 34(9), 942–949. 10.1038/nbt.3601

Grasset, C., Mendonça, R., Villamor Saucedo, G., Bastviken, D., Roland, F., & Sobek, S. (2018). Large but variable methane production in anoxic freshwater sediment upon addition of allochthonous and autochthonous organic matter. Limnology and Oceanography, 63(4), 1488–1501. 10.1002/lno.10786

Grün, B., & Hornik, K. (2011). **topicmodels**: An *R* Package for Fitting Topic Models. Journal of Statistical Software, 40(13). 10.18637/jss.v040.i13

Guillemette, F., von Wachenfeldt, E., Kothawala, D. N., Bastviken, D., & Tranvik, L. J. (2017). Preferential sequestration of terrestrial organic matter in boreal lake sediments. Journal of Geophysical Research: Biogeosciences, 122(4), 863–874. 10.1002/2016JG003735

Heathcote, A. J., Anderson, N. J., Prairie, Y. T., Engstrom, D. R., & del Giorgio, P. A. (2015). Large increases in carbon burial in northern lakes during the Anthropocene. Nature Communications, 6(1), Article 1. 10.1038/ncomms10016

Holgerson, M. A., & Raymond, P. A. (2016). Large contribution to inland water CO2 and CH4 emissions from very small ponds. Nature Geoscience, 9(3), Article 3. 10.1038/ngeo2654

Hornbach, D. J., Shea, K. L., Dosch, J. J., Thomas, C. L., Gartner, T. B., Aguilera, A. G., Anderson, L. J., Geedey, K., Mankiewicz, C., Pohlad, B. R., & Schultz, R. E. (2021). Decomposition of Leaf Litter from Native and Nonnative Woody Plants in Terrestrial and Aquatic Systems in the Eastern and Upper Midwestern U.S.A. The American Midland Naturalist, 186(1), 51–75. 10.1674/0003-0031-186.1.51

Intergovernmental Panel On Climate Change (Ipcc). (2023). Climate Change 2022 – Impacts, Adaptation and Vulnerability: Working Group II Contribution to the Sixth Assessment Report of the Intergovernmental Panel on Climate Change (1st ed.). Cambridge University Press. 10.1017/9781009325844

Karlsson, J., Berggren, M., Ask, J., Byström, P., Jonsson, A., Laudon, H., & Jansson, M. (2012). Terrestrial organic matter support of lake food webs: Evidence from lake metabolism and stable hydrogen isotopes of consumers. Limnology and Oceanography, 57(4), 1042–1048. 10.4319/lo.2012.57.4.1042

Karlsson, J., Byström, P., Ask, J., Ask, P., Persson, L., & Jansson, M. (2009). Light limitation of nutrient-poor lake ecosystems. Nature, 460(7254), Article 7254. 10.1038/nature08179

Kolde, R. (2019). Pheatmap: Pretty Heatmaps. R package version 1.0.12. (Version v.1.0.12) [Computer software]. https://CRAN.R-project.org/package=pheatmap

Kozich, J. J., Westcott, S. L., Baxter, N. T., Highlander, S. K., & Schloss, P. D. (2013). Development of a dual-index sequencing strategy and curation pipeline for analyzing amplicon sequence data on the MiSeq Illumina sequencing platform. Applied and Environmental Microbiology, 79(17), 5112– 5120. 10.1128/AEM.01043-13

Kudo, H., Cheng, K.-J., & Costerton, J. W. (1987). Interactions between Treponema bryantii and cellulolytic bacteria in the in vitro degradation of straw cellulose. Canadian Journal of Microbiology, 33(3), 244–248. 10.1139/m87-041

Kurade, M. B., Saha, S., Salama, E.-S., Patil, S. M., Govindwar, S. P., & Jeon, B.-H. (2019). Acetoclastic methanogenesis led by Methanosarcina in anaerobic co-digestion of fats, oil and grease for enhanced production of methane. Bioresource Technology, 272, 351–359. 10.1016/j.biortech.2018.10.047

Lapierre, J.-F., Guillemette, F., Berggren, M., & del Giorgio, P. A. (2013). Increases in terrestrially derived carbon stimulate organic carbon processing and CO2 emissions in boreal aquatic ecosystems. Nature Communications, 4(1), Article 1. 10.1038/ncomms3972

Leadbeater, D. R., Oates, N. C., Bennett, J. P., Li, Y., Dowle, A. A., Taylor, J. D., Alponti, J. S., Setchfield, A. T., Alessi, A. M., Helgason, T., McQueen-Mason, S. J., & Bruce, N. C. (2021). Mechanistic strategies of microbial communities regulating lignocellulose deconstruction in a UK salt marsh. Microbiome, 9, 48. 10.1186/s40168-020-00964-0

Love, M., Ahlmann-Eltze, C., Forbes, K., Anders, S., & Huber, W. (2020). *DESeq2: Differential gene expression analysis based on the negative binomial distribution* (Version 1.30.0) [Computer software]. Bioconductor version: Release (3.12). 10.18129/B9.bioc.DESeq2

Lyu, Z., Shao, N., Akinyemi, T., & Whitman, W. B. (2018). Methanogenesis. Current Biology, 28(13), R727–R732. 10.1016/j.cub.2018.05.021

Mand, T. D., & Metcalf, W. W. (2019). Energy Conservation and Hydrogenase Function in Methanogenic Archaea, in Particular the Genus Methanosarcina. Microbiology and Molecular Biology Reviews: MMBR, 83(4), e00020–19. 10.1128/MMBR.00020-19

McMurdie, P. J., & Holmes, S. (2013). phyloseq: An R Package for Reproducible Interactive Analysis and Graphics of Microbiome Census Data. PLOS ONE, 8(4), e61217. 10.1371/journal.pone.0061217

Nemergut, D. R., Costello, E. K., Hamady, M., Lozupone, C., Jiang, L., Schmidt, S. K., Fierer, N., Townsend, A. R., Cleveland, C. C., Stanish, L., & Knight, R. (2011). Global patterns in the biogeography of bacterial taxa. Environmental Microbiology, 13(1), 135–144. 10.1111/j.1462-2920.2010.02315.x

Nikita, M. (2020). ldatuning: Tuning of the Latent Dirichlet Allocation Models Parameters (Version 1.0.2) [Computer software]. https://CRAN.R-project.org/package=ldatunin

Niño-García, J. P., Ruiz-González, C., & del Giorgio, P. A. (2016). Interactions between hydrology and water chemistry shape bacterioplankton biogeography across boreal freshwater networks. The ISME Journal, 10(7), 1755–1766. 10.1038/ismej.2015.226

Oksanen, J., Kindt, R., Legendre, P., Hara, B., Simpson, G., Solymos, P., Henry, M., Stevens, H., Maintainer, H., & Oksanen@oulu, jari. (2009). The vegan Package.

Ou, Y.-F., Dong, H.-P., McIlroy, S. J., Crowe, S. A., Hallam, S. J., Han, P., Kallmeyer, J., Simister, R. L., Vuillemin, A., Leu, A. O., Liu, Z., Zheng, Y.-L., Sun, Q.-L., Liu, M., Tyson, G. W., & Hou, L.-J. (2022). Expanding the phylogenetic distribution of cytochrome b-containing methanogenic archaea sheds light on the evolution of methanogenesis. The ISME Journal, 16(10), Article 10. 10.1038/s41396-022-01281-0

Pace, M. L., Cole, J. J., Carpenter, S. R., Kitchell, J. F., Hodgson, J. R., Van de Bogert, M. C., Bade, D. L., Kritzberg, E. S., & Bastviken, D. (2004). Whole-lake carbon-13 additions reveal terrestrial support of aquatic food webs. Nature, 427(6971), Article 6971. 10.1038/nature02227

Polis, G. A., Anderson, W. B., & Holt, R. D. (1997). Toward an Integration of Landscape and Food Web Ecology: The Dynamics of Spatially Subsidized Food Webs. Annual Review of Ecology and Systematics, 28(1), 289–316. 10.1146/annurev.ecolsys.28.1.289

Quast, C., Pruesse, E., Yilmaz, P., Gerken, J., Schweer, T., Yarza, P., Peplies, J., & Glöckner, F. O. (2013). The SILVA ribosomal RNA gene database project: Improved data processing and web-based tools. Nucleic Acids Research, 41(Database issue), D590–D596. 10.1093/nar/gks1219

R Core Team. (2018). R: A language and environment for statistical computing [Computer software]. R Foundation for Statistical Computing. https://www.R-project.org/

Ruuskanen, M. O., St. Pierre, K. A., St. Louis, V. L., Aris-Brosou, S., & Poulain, A. J. (2018). Physicochemical Drivers of Microbial Community Structure in Sediments of Lake Hazen, Nunavut, Canada. Frontiers in Microbiology, 9. https://www.frontiersin.org/articles/10.3389/fmicb.2018.01138

Saha, S., Kurade, M. B., Ha, G.-S., Lee, S. S., Roh, H.-S., Park, Y.-K., & Jeon, B.-H. (2021). Syntrophic metabolism facilitates Methanosarcina-led methanation in the anaerobic digestion of lipidic slaughterhouse waste. Bioresource Technology, 335, 125250. 10.1016/j.biortech.2021.125250

Salama, E.-S., Saha, S., Kurade, M. B., Dev, S., Chang, S. W., & Jeon, B.-H. (2019). Recent trends in anaerobic co-digestion: Fat, oil, and grease (FOG) for enhanced biomethanation. Progress in Energy and Combustion Science, 70, 22–42. 10.1016/j.pecs.2018.08.002

Sankaran, K., & Holmes, S. P. (2019). Latent variable modeling for the microbiome. Biostatistics, 20(4), 599–614. 10.1093/biostatistics/kxy018

Schloss, P. D., Westcott, S. L., Ryabin, T., Hall, J. R., Hartmann, M., Hollister, E. B., Lesniewski, R. A., Oakley, B. B., Parks, D. H., Robinson, C. J., Sahl, J. W., Stres, B., Thallinger, G. G., Van Horn, D. J., & Weber, C. F. (2009). Introducing mothur: Open-Source, Platform-Independent, Community-Supported Software for Describing and Comparing Microbial Communities. Applied and Environmental Microbiology, 75(23), 7537–7541. 10.1128/AEM.01541-09

Sobek, S., Durisch-Kaiser, E., Zurbrügg, R., Wongfun, N., Wessels, M., Pasche, N., & Wehrli, B. (2009). Organic carbon burial efficiency in lake sediments controlled by oxygen exposure time and sediment source. Limnology and Oceanography, 54(6), 2243–2254. 10.4319/lo.2009.54.6.2243

Song, N., Xu, H., Yan, Z., Yang, T., Wang, C., & Jiang, H.-L. (2019). Improved lignin degradation through distinct microbial community in subsurface sediments of one eutrophic lake. Renewable Energy, 138, 861–869. 10.1016/j.renene.2019.01.121

Stackebrandt, E., & Goebel, B. M. (1994). Taxonomic Note: A Place for DNA-DNA Reassociation and 16S rRNA Sequence Analysis in the Present Species Definition in Bacteriology. International Journal of Systematic and Evolutionary Microbiology, 44(4), 846–849. 10.1099/00207713-44-4-846

Stanley, E. H., Casson, N. J., Christel, S. T., Crawford, J. T., Loken, L. C., & Oliver, S. K. (2016). The ecology of methane in streams and rivers: Patterns, controls, and global significance. Ecological Monographs, 86(2), 146–171. 10.1890/15-1027

Tardy, V., Etienne, D., Millet, L., & Lyautey, E. (2022). Lake Sediments From Littoral and Profundal Zones are Heterogeneous but Equivalent Sources of Methane Produced by Distinct Methanogenic Communities—A Case Study From Lake Remoray. Journal of Geophysical Research: Biogeosciences, 127(11), e2021JG006776. 10.1029/2021JG006776

Thauer, R. K., Kaster, A.-K., Seedorf, H., Buckel, W., & Hedderich, R. (2008). Methanogenic archaea: Ecologically relevant differences in energy conservation. Nature Reviews Microbiology, 6(8), Article 8. 10.1038/nrmicro1931

Tittel, J., Hüls, M., & Koschorreck, M. (2019). Terrestrial Vegetation Drives Methane Production in the Sediments of two German Reservoirs. Scientific Reports, 9(1), Article 1. 10.1038/s41598-019-52288-1

Vanwonterghem, I., Evans, P. N., Parks, D. H., Jensen, P. D., Woodcroft, B. J., Hugenholtz, P., & Tyson, G. W. (2016). Methylotrophic methanogenesis discovered in the archaeal phylum Verstraetearchaeota. Nature Microbiology, 1(12), Article 12. 10.1038/nmicrobiol.2016.170

Vincent, W., Kumagai, M., & Couture, R.-M. (2023). Wetzel’s Limnology—Sediments and Microbiomes (4th ed.).

Walter E. Dean, Jr. (1974). Determination of Carbonate and Organic Matter in Calcareous Sediments and Sedimentary Rocks by Loss on Ignition: Comparison With Other Methods. SEPM Journal of Sedimentary Research, *Vol.* 44. 10.1306/74D729D2-2B21-11D7-8648000102C1865D

Ward, T. E., & Frea, J. I. (1980). Sediment Distribution of Methanogenic Bacteria in Lake Erie and Cleveland Harbor. Applied and Environmental Microbiology, 39(3), 597–603. 10.1128/aem.39.3.597-603.1980

West, W. E., Coloso, J. J., & Jones, S. E. (2012). Effects of algal and terrestrial carbon on methane production rates and methanogen community structure in a temperate lake sediment. Freshwater Biology, 57(5), 949–955. 10.1111/j.1365-2427.2012.02755.x

West, W. E., Creamer, K. P., & Jones, S. E. (2016). Productivity and depth regulate lake contributions to atmospheric methane. Limnology and Oceanography, 61(S1), S51–S61. 10.1002/lno.10247

Wilkinson, G. M., Pace, M. L., & Cole, J. J. (2013). Terrestrial dominance of organic matter in north temperate lakes. Global Biogeochemical Cycles, 27(1), 43–51. 10.1029/2012GB004453

Winkel, M., Trivedi, C. B., Mourot, R., Bradley, J. A., Vieth-Hillebrand, A., & Benning, L. G. (2022). Seasonality of Glacial Snow and Ice Microbial Communities. Frontiers in Microbiology, 13. https://www.frontiersin.org/articles/10.3389/fmicb.2022.876848

Yakimovich, K. M., Orland, C., Emilson, E. J. S., Tanentzap, A. J., Basiliko, N., & Mykytczuk, N. C. S. (2020). Lake characteristics influence how methanogens in littoral sediments respond to terrestrial litter inputs. The ISME Journal. 10.1038/s41396-020-0680-9

Yu, Y., Lee, C., Kim, J., & Hwang, S. (2005). Group-specific primer and probe sets to detect methanogenic communities using quantitative real-time polymerase chain reaction. Biotechnology and Bioengineering, 89(6), 670–679. 10.1002/bit.20347

Zhang, C. (2022). MicrobiomeStat: Statistical Methods for Microbiome Compositional Data. (Version 1.1) [Computer software]. <https://CRAN.R-project.org/package=MicrobiomeStat>.

Zhang, J., Mao, L., Zhang, L., Loh, K.-C., Dai, Y., & Tong, Y. W. (2017). Metagenomic insight into the microbial networks and metabolic mechanism in anaerobic digesters for food waste by incorporating activated carbon. Scientific Reports, 7(1), Article 1. 10.1038/s41598-017-11826-5

Zhang, J., Yang, Y., Zhao, L., Li, Y., Xie, S., & Liu, Y. (2015). Distribution of sediment bacterial and archaeal communities in plateau freshwater lakes. Applied Microbiology and Biotechnology, 99(7), 3291–3302. 10.1007/s00253-014-6262-x

Zhou, H., He, K., Chen, J., & Zhang, X. (2022). LinDA: linear models for differential abundance analysis of microbiome compositional data. arXiv. https://arxiv.org/abs/2104.00242

